# Confinement-induced transition between wave-like collective cell migration modes

**DOI:** 10.1101/495747

**Authors:** Vanni Petrolli, Magali Le Goff, Monika Tadrous, Kirsten Martens, Cédric Allier, Ondrej Mandula, Lionel Hervé, Silke Henkes, Rastko Sknepnek, Thomas Boudou, Giovanni Cappello, Martial Balland

## Abstract

The structural and functional organization of biological tissues relies on the intricate interplay between chemical and mechanical signaling. Whereas the role of constant and transient mechanical perturbations is generally accepted, several studies recently highlighted the existence of long-range mechanical excitations (i.e., waves) at the supracellular level. Here, we confine epithelial cell mono-layers to quasi-one dimensional geometries, to force the establishment of tissue-level waves of well-defined wavelength and period. Numerical simulations based on a self-propelled Voronoi model reproduce the observed waves and exhibit a phase transition between a global and a multi-nodal wave, controlled by the confinement size. We confirm experimentally the existence of such a phase transition, and show that wavelength and period are independent of the confinement length. Together, these results demonstrate the intrinsic origin of tissue oscillations, which could provide cells with a mechanism to accurately measure distances at the supracellular level.

---

Supracellular organization plays a key role in establishing and maintaining structure, function and homeostasis in tissues. In the early stages of embryonic development, where features need to arise spontaneously from a homogeneous state, this organization closely follows morphogenic chemical patterns. Turing pioneered the idea that symmetry breaking results from linear instabilities in the reaction-diffusion dynamics of morphogens [1]. In the most general case, however, chemical reactions, osmotic pressures and mechanical forces all cooperate to determine the tissue-level organization. This is confirmed by an increasing number of recent studies, which indicate that cell proliferation, differentiation and motility are strongly impacted by the physical properties of the microenvironment [2–6]. Therefore, a full physical understanding of tissue mechanics and morphogenesis requires treating chemical and mechanical effects simultaneously. Several recent works reported that wave-like patterns of the local cell velocity spontaneously appear in colonies of epithelial cells [7, 8]. Those velocity waves have also been observed in spreading epithelial sheets [9–11], regardless of cell proliferation [12], and are correlated to oscillations of the forces exerted by the cells on the substrate [13]. Such long wavelength patterns also appear in confined geometries where cell migration is limited to local cell rearrangements [14–18]. These waves are characterized by a wavelength λ and a period *T*, and show a surprisingly large spatial and temporal coherence. They can be modelled either at the particle level [17] or using continuum approaches [12, 18], based on a coupling between cell motility and intercellular forces. In this Letter, we explore whether period and wavelength of collective wave excitations in epithelial cell monolayers are intrinsically encoded in the activity of the cell, or if they are affected by external constraints such as a specific set of boundary conditions. To achieve this, we analyzed the collective motion of epithelial cells confined to a quasi-one-dimensional channel. The experiments were accompanied by a series of numerical simulations, based on a self-propelled Voronoi model (SPV) [19–21], adapted to take into account the confining geometry. Our results show that tuning the length of the confining channel drives a phase transition between a state of global oscillations and a multinodal wave state. This transition is a consequence of the interplay between local cell active dynamics and global confinement. The effect is robust and does not require detailed knowledge of molecular processes but relies on a simple polarity-velocity alignment mechanism studied in the physics of dense active matter systems.

To confine cells to a quasi one-dimensional pattern, we prepared adherent stripes on soft polyacrylamide gels (E ≃ 40 kPa), as described previously [22] (outlined in Fig. 1a). Stripes of different length (100 to 2000 *μ*m), but of the same width (40 *μ*m), were patterned on the same substrate. Epithelial Madin-Darby Canine Kidney (MDCK) cells were then seeded on the patterned substrates with initial concentration of 2.5 ± 0.5×10^4^ cells/cm^2^. The samples were washed with fresh medium 1h after seeding, then placed in the incubator (37° C and 5% CO_2_) until the end of the experiments. Cells were imaged *in-situ* unsing in-line holographic (defocus) microscope (see SI and Fig SI-1) [23] for ≃ 48 hours after confluence, gathering one image every 10 minutes (e.g., Fig. 1a-Phase). Cell velocities were computed with a custom-made Particle Image Velocimetry algorithm (PIV) with a final resolution of 20 min and 14 *μ*m. To generate the kymograph, we cropped the videos in time to consider only confluent tissues, in an interval where the average absolute velocity was higher than 4 *μ*m/h [24]. We then averaged the horizontal component of the speed along the transverse direction *υ_x_* (*x; t*) = (*υ* (*x,y; t*))_*y*_. Wex removed low frequency drifts using a Gaussian high-pass filter cropping 50% of the signal at 700 *μm* and 10 h. The kymograph in Fig. 1b represents the spatio-temporal evolution of the velocity field over 22 hours and over the whole stripe. Fig. 1c shows the instantaneous velocity profile, where a periodic oscillation of the speed appears. To quantify the period and the wavelength of these oscillations, we computed the autocorrelation function of the kymograph *g*(*δx,δt*) = 〈*υ_x_* (*x,t*) *υ_x_* (*x + δx,t* + *δt*))_*x,t*_, displayed in Fig. 1d. We observe the appearance of an oscillating pattern in the autocorrelation function, both along the spatial and the temporal directions (respectively shown in panels e and f in Fig. 1). This pattern indicates the establishment of an extended multi-nodal standing wave, with the wavelength and period λ = 370 ± 30μm and T = 4.7 ± 0.7 h, respectively (errors represent the standard deviation, n = 59) (see histograms in Fig. 1e-f).

**FIG. 1.**
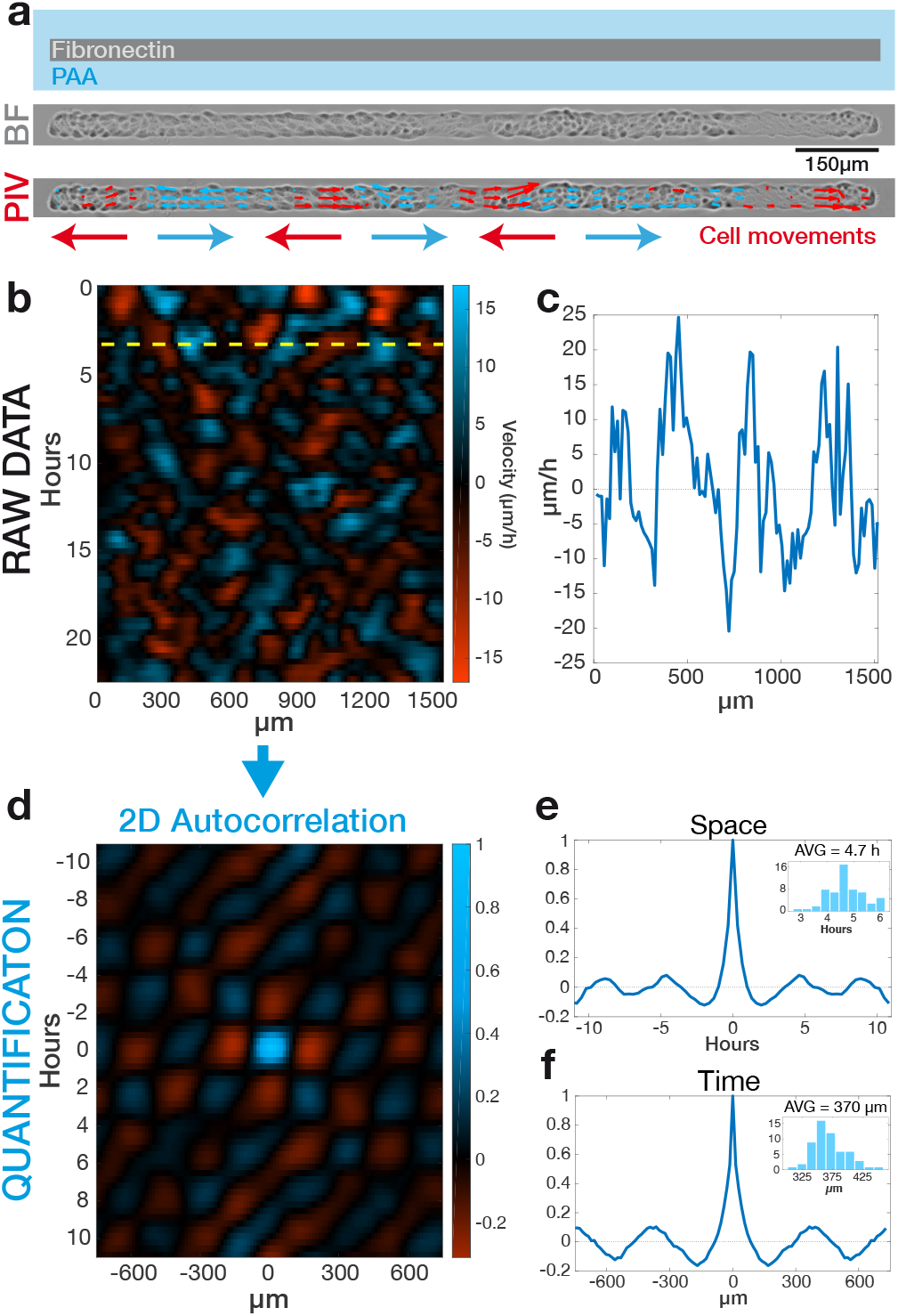
(a) Top: MDCK cells are seeded onto a polyacrylamide (PA) gel patterned with fibronectin stripes (width: 40 *μ*m, length: 100-2000 *μ*m). Center: Phase-contrast image of confluent tissue confined to a 1500 *μ*m long stripe and the relative velocity field measured by PIV (bottom). Velocities pointing in the positive (negative) x-axis direction are shown in blue (red), in agreement with the arrows reported under the image. b) Kymograph representing the average horizontal velocity *υ_||_* (*x; t*) in time and and an example of velocity profile (c) along the dashed line. We removed low frequency drifts using a Gaussian high-pass filter. To quantify the periodicity of oscillations, we calculate the spatio-temporal autocorrelation of the kymograph (d) and measure peak spacing along the spatial (e) and temporal coordinates (f) (insets: distribution of peak periodicity for n=59 independent stripes). Images in panels (d) and (f) were smoothed for visualization purposes with a low-pass Gaussian filter (*σ_x_*=15 *μ*m, *σ_t_*=30 min). Smoothing was not applied to panels (e), (g) and (h).

To obtain a detailed understanding of oscillations in tissues, we consider a computational framework based on a recently introduced self-propelled Voronoi model (SPV) [19–21]. The model used in this study is similar to that used in Ref. [21] to describe flocking transitions in confluent tissues, with an important modification that instead of using periodic boundary conditions, we imposed confinement through a repulsive rectangular wall of size (*L_x_*, *L_y_*), to reproduce the experiments geometry. Full details of the model and its implementation can be found in Ref. [20] (also see Supplemental Material for the parameters used). Briefly, the confluent cell monolayer is modeled as a two-dimensional network of Voronoi polygons covering the plane (Voronoi tessellation of all cell centre positions, see Fig. 2a). Each configuration of cells is described by the positions of cell centroids with its energy given by the commonly used Vertex Model [25], which depends on the area and perimeter of each cell. The parameters of the Vertex model include area and perimeter stiffness constants (*K* and *Γ*) and target area and perimeter (*A*^0^ and *P*^0^). These parameters were chosen to describe a monolayer in a solid like regime (with a shape factor 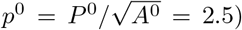) [19, 26], to avoid shear flows induced by the boundaries. As in Refs. [19–21], we consider an overdamped dynamics, i.e., a force balance between frictional force with the substrate, self-propulsion at a constant velocity *v***0** along the direction of cell polarity, **n**_*i*_, and mechanical forces between the cells determined as a negative gradient with respect to cell position of the SPV model energy functional. The value of *v***0** can be set to match the experimental observations (Fig. 2b-d), but does not affect the general oscillatory behavior. The dynamics of the cell polarity **n**_*i*_, described by the angle *θ_i_* with the x-axis of the laboratory reference frame (i.e., **n**_*i*_ = (cos(*θ_i_*), sin(*θ_i_*))) is

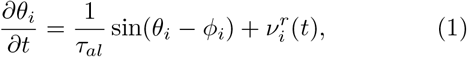

with *ϕ_i_* being the angle between the velocity of cell *i* and the x-axis, and 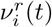 being an orientational Gaussian noise. The angular dynamics is thus controlled by the interplay of rotational diffusion (kept constant in this study) and the polarity-velocity alignment with rate 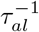, with *τ_al_* being the time required by the cell to reorient its polarization in the direction of its velocity. This feedback mechanism leads to oscillations in confinement, where *τ_al_* plays the role of an effective inertia, and the oscillations are along the lowest-energy elastic modes of the material [27]. This feedback mechanism is also at the origin of flocks of in non-confined tissues [21].

**FIG. 2.**
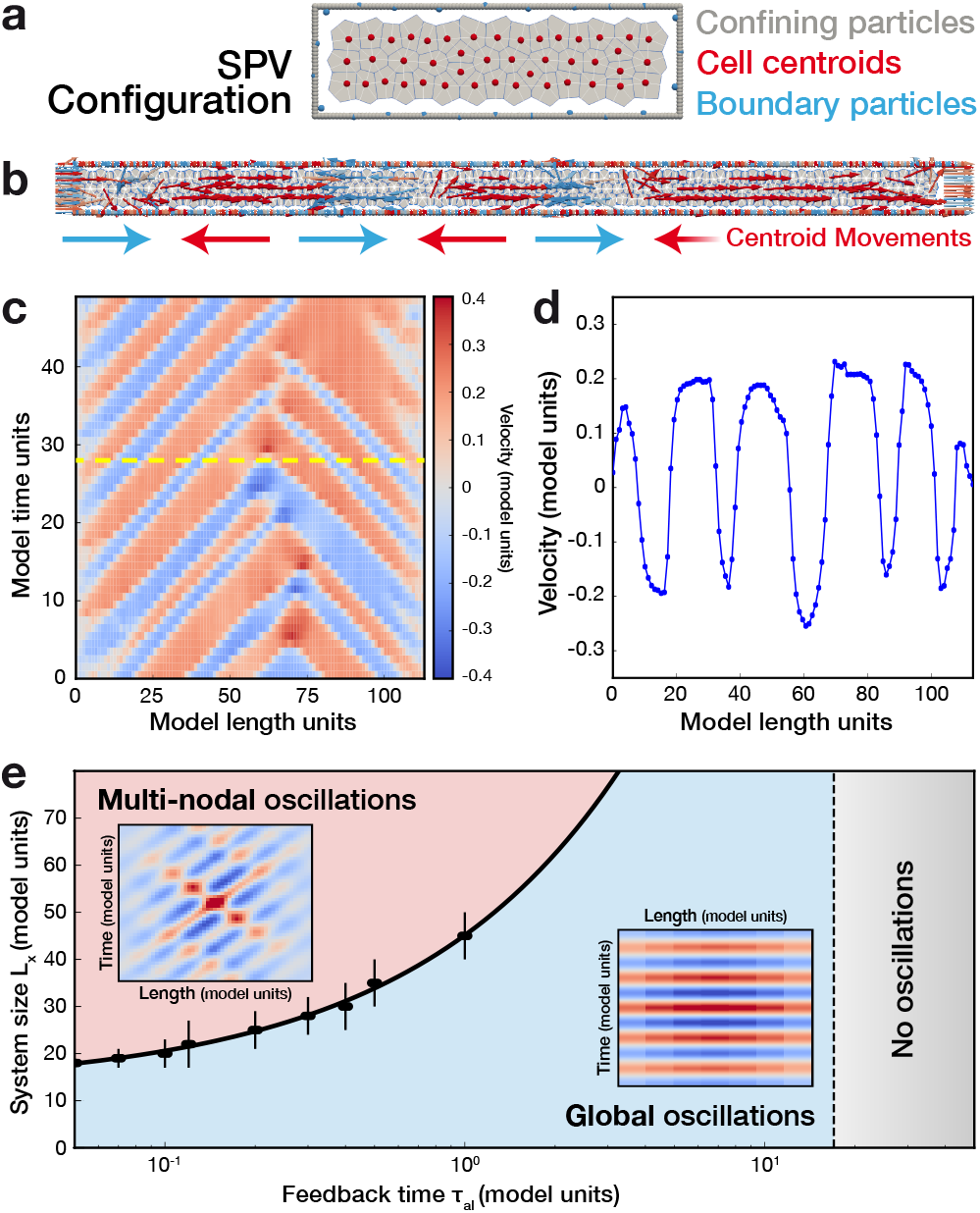
Self-propelled Voronoi Model for collective oscillations in confluent tissues. (a) Example of tissue configuration obtained from the integration of the SPV model. Voronoi tessellation of the plane and centroid positions. (b) Velocity field of the centroids of the tesselation. Velocities pointing to the positive direction on the x-axis are represented in red and to the negative direction in blue. (c) Kymograph representing the average horizontal velocity (*υ*_||_ (*x; t*)) over time and its profile (d) along the dotted line. (e) Phase diagram of oscillation patterns in the monolayer. Oscillations are observed for small enough values of the feedback timescale (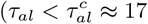 model time units). For *τ_al_* < *τ_al_*, multi-nodal oscillations are observed for large systems *L_Y_* > *L_c_* (*τ_al_*) whereas small systems oscillate as a block (global oscillations). The dots delimiting the two phases indicate the value of *L_c_*(*τ_al_*) and are determined by the number of nodes in the spatial velocity autocorrelation functions (0 in the case of global oscillations) and the full line is a power law fit. Insets show the autocorrelation of the typical kymographs of x-velocity in the corresponding phases.

Simulations of confined tissue layers show steady state oscillations akin to those observed in experiments (Fig. 2b) In the following, we study the dependence of these oscillations on the confining length *L_X_* and show that a feedback mechanism for alignment (through *τ_al_*) is key to observe such mechanical waves in SPV. First we consider the case of long confining channels, where multinodal oscillations were observed in the experiments (Fig. 1). The simulation results displayed in Fig. 2c-d are obtained for a system with the same confining length (about 3 cells in *y*—direction) and aspect ratio as in the experiments in Fig. 1. We observe a pattern in the x-component of the velocity, *υ_x_*, and using the same analysis tools as in Fig. 1, we extract the wavelength *λ_SPV_* ≈ 22 model length units and the period *T_SPV_* ≈ 8 model time units. Note that by approximately matching the timescale of the model to the experiments (through the cells velocity), one would get from these simulation data *λ* ≈ 300 *μ*m and for the period *T* ≈ 2 hours. This indicates that this model is able to reproduce the features observed in the experiments, although some fine tuning of parameters is required for a quantitative match. Note that although the instantaneous velocity profiles and autocorrelation plots appear to be similar to the experiments, the full spatio-temporal dynamics of the model does not corresponds to standing wave oscillations. If the system size *L_X_* is decreased (keeping the value of *τ_al_* constant), the number of nodes also decreases up to a point where the system size can only accommodate a single spatial period of oscillation, and the system reaches a regime of global oscillation, where the direction of motion of all cells is coordinated. This transition is illustrated in Fig. 2e. The feedback timescale also plays an important role as no oscillations are observed if *τ_al_* is too large (i.e., the noise dominates over the coupling), and the critical length at which one observes multi-nodal oscillations increases with *τ_al_*. In the small system regime, the period of oscillations increases linearly with the system size as previously reported [17, 18], and also with *τ_al_* (until oscillations eventually vanish for large values of *τ_al_*), consistent with the role of the feedback mechanism as an effective inertia [27]. Therefore, the SPV model describes the transition controlled by the stripe length *L_c_* (*τ_al_*) between global oscillations where all cells coordinate their motion to a regime where patches of cells coordinate their motion direction locally.

To verify this prediction, we varied the length *L_X_* of the stripe between 100 *μ*m and 2000 *μ*m (Fig. 3a), in order to tune the system across the critical length *L_c_*. In approximately 95% of experiments, in agreement with model predictions, we observed two types of behaviors: 1) A global movement of all cells that alternates between rightward and leftward motion (as seen from the autocorrelation function of the kymograph in Fig. 3b) and 2) The establishment of a multinodal standing wave with the antinodal cells moving back and forth, while cells in the nodes are being alternately compressed and dilated.

**FIG. 3.**
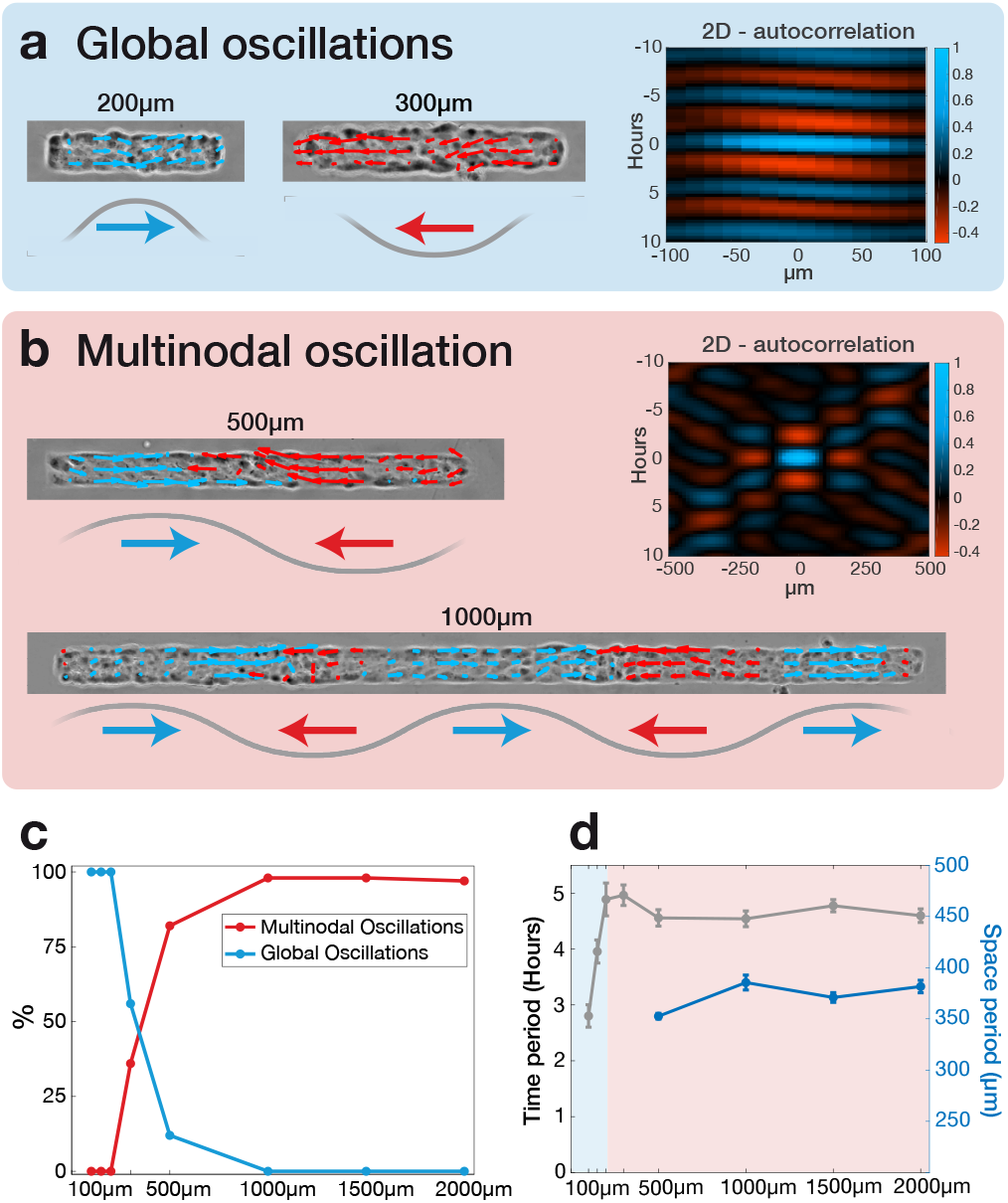
Dependence of oscillatory behaviour on the stripe length. a) The velocity field superimposed on phase contrast images for short stripes of length 200 *μ*m and 300 *μ*m displays global oscillations, generating a characteristic twodimensional autocorrelation (right). Longer lines (500 *μ*m and 1000 *μ*m) display multinodal oscillations (b), which give rise a different pattern in the autocorrelation image. Velocities pointing in the positive x-axis direction are represented in blue, those pointing in the negative x-axis direction are represented in red, in agreement with the arrows reported in the schemes under each image. For each length, we display the frequency of each phenotype (c) and the characteristic time and space periodicity (d) calculated. Bars represent the standard error of the mean.

The incidence of the two behaviors strongly depends on *L*, with a transition for *L* ≃ *λ*. In the experiments with *L* < 200 *μ*m, the global oscillation statistically dominated. In this case, the period scales linearly with the tissue size (Fig. 3d, blue area), while the wavelength is imposed by the confinement. In large structures (*L* > 500 *μ*m), we only found multinodal waves, with the period and wavelength independent of *L* (Fig. 3d, red area). Fig. 3c quantifies the transition, with on average 39 tissues per point, obtained from three independent experiments. Our experiments confirmed the existence of a self-sustained oscillatory mode in the epithelial layers. Using the typical period and wavelength, we can define an effective velocity *u_υ_* = *L/T* ≃ 78 ± 13 *μ*m/h, which is independent of the pattern size. Even for small patterns (*L* <500 *μ*m), this velocity is preserved as the period scales linearly with the pattern length. We also note that *u_φ_* is approximately ten-fold larger than the average speed of individual cells within the epithelial layer (between 4 and 12 *μ*m/h, depending on cell density [24, 28]). Eventually, the spatial coherence of supracellular waves exceeds the largest pattern observable with our microscope.

Simulations using the SPV model show the emergence of sustained collective oscillations in confined monolayers. We identified two crucial conditions to produce these oscillations: 1) The existence of a delayed feedback between the cell velocity and self-propulsion direction that introduces a new timescale in the dynamics and 2) A very limited number of cellular rearrangement, at the limit of the solid-like regime. These ingredients allow the system to be described by linear elasticity, and for oscillations along the lowest energy elastic modes to dominate the dynamics [27]. One could thus envision tuning the oscillations by controlling cell-cell interactions through RAB5 or cadherin-mediated junctions, without affecting cells’ individual mobility [21, 29]. Contrarily to experiments, where multi-nodal standing waves are observed, the SPV model describes propagating oscillations. Several reasons may explain the difference. First, a standing wave is established only when the wavelength exactly matches the boundary conditions. Thus, a fine-tuning of the pattern length is needed in models, while the intrinsic variability between cells could make the real epithelium more adaptable to small variations of the confinement size. Second, a different choice of the coupling mechanism could also introduce a new timescale in the model and better describe standing waves in confined tissue. Two-dimensional SPV models are usually adapted to describe spatially extended monolayers, while the stringent confinement used experimentally makes the system be quasi-onedimensional. Moreover, whereas in the experiments the cells appear to be more elongated near the boundary, the constraint of maintaining a Delaunay triangulation (dual of the Voronoi tesselation) in the SPV prevents elongated cell configurations, and hence the tissue configuration in simulation do not realistically capture the shape of cells near the containment boundary. In conclusion, we demonstrate that the typical period and wavelength of epithelial tissue oscillations are intrinsically encoded in the cells, and are not adapted to external confinements. For this system, our SPV model predicts a transition between global oscillation and multi-nodal waves, the existence of which is confirmed experimentally for a pattern length *L_c_* ≃ 400 *μ*m. From a biological perspective this transition could be significant. If in small systems all the cells behave similarly - the entire layer alternately moves back and forth - in large systems cells located either in the nodes or in the anti-nodes experience different mechanical stimuli and may undergo different fates, which can ultimately lead to supracellular patterning. The existence of an intrinsic wavelength λ also provides an intrinsic metric, likely encoded in the cell. It is interesting to note that λ roughly corresponds to the typical sizes of an embryo (e.g. in Drosophyla embryo, both length and circumference approaches 400-500 *μ*m). Based on this, two important biological questions arise. Is this intrinsic metric used by the organism to measure distance inside a developing embryo? Does a collective long range excitation allow cells to probe their distant environment, in a timescale much shorter that that allowed by their own motility?

The authors would like to acknowledge I. Wang for the development of Particle Image Velocimetry and and P. Moreau for his technical support. We also thank K. Hennig, T. Andersen and C. Guilluy for valuable and supportive discussions. GC has been supported by the Institut National de la Sante et de la Recherche Medicale (Grant ‘‘Physique et Cancer” PC201407). T.B. and M.B. acknowledge financial support from the CNRS “Mission pour l’lnterdisciplinarité” and the Center of Excellence of Multifunctional Architectured Materials “CEMAM” (n AN-10-LABX-44-01). M.B. acknowledges financial support from the ANR MechanoSwitch project, grant ANR-17-CE30-0032-01 of the French Agence Nationale de la Recherche. RS and SH the UK BBSRC, award numbers BB/N009789/1 and BB/N009150/1-2.

