## Supplementary material for "Confinement-induced transition between wave-like collective cell migration modes"

### Supplemental material for:
### "Confinement-induced transition between wave-like collective cell migration modes"

Vanni Petrolli,[1] Magali Le Goff,[1] Monika Tadrous,[1] Kirsten Martens,[1] Cédric Allier,[2] Ondrej Mandula,[2] Lionel Hervé,[2] Silke Henkes,[3] Rastko Sknepnek,[4,5] Thomas Boudou,[6] Giovanni Cappello,[6,*] and Martial Balland[6,†]

[1] *Université Grenoble Alpes, Laboratoire Interdisciplinaire de Physique, CNRS, F-38000 Grenoble, France*
[2] *CEA, LETI, MINATEC, 17 rue des Martyrs, 38054 Grenoble cedex 9, France*
[3] *Department of Mathematics, University of Bristol, Bristol BS81TW, United Kingdom*
[4] *School of Science and Engineering, University of Dundee, Dundee DD1 4HN, United Kingdom*
[5] *School of Life Sciences, University of Dundee, Dundee DD1 5EH, United Kingdom*
[6] *Université Grenoble Alpes, Laboratoire Interdisciplinaire de Physique, CNRS, F-38000 Grenoble, France*
(Dated: December 13, 2018)

#### PHASE FROM DEFOCUS MICROSCOPY

**Experimental Setup** We used low-cost components to build a holographic video-microscope in inverted configuration. The setup and a schematic drawing is shown Fig. 1. Blue LED (CREE, max 450 nm, FWHM 18 nm) source coupled with 400 $\mu$m multimode fibre (Thorlabs) with a narrow band filter (Thorlabs FB450-10, max 450 nm, FWHM 10 nm) is used for illumination in transmission geometry. Semi-coherent light passes through the sample and is collected by 10x/0.25 NA Objective (Motic CCIS EF-N Plan, Achromat). A short tube lens (Thorlabs, AC254-050-A, $f$ = 50 mm) is used to create an image on a CMOS sensor (IDS UI-1492LE). Standard Thorlabs components are used for housing of the optics and the camera. Image acquisition with synchronized illumination is controlled with Raspberry Pi[1]. Note, that the reconstruction of the data is performed on a different computer. The field of view of our system with 10x/0.25NA objective is $2.3 \times 1.6 = 3.7$ mm$^2$ and the spatial resolution is approximately $3\mu m$. With the current camera and Raspberry Pi we can perform time-laps movies with time resolution of several seconds.

**Data Acquisition** Data are acquired approximately 50 $\mu$m out-of-focus by physically moving either the sample or the objective. The out-of-focus distance does not require high precision as the correct defocus value is determined post-acquisition. Note that there is no mechanical movement of the sample or the objective during the time-laps acquisition. The defocus is performed before the start of the measurement and left for the rest of the experiment. The potential slight drift in the axial direction can be compensated in the reconstruction process post-acquisition, however, a rigid construction of our microscope reduces the axial drift and vibrations to negligible levels.

**Data Reconstruction** Our reconstruction algorithm is based on an iterative optimization of Fresnel diffraction model for coherent light[2]. The fact, that the illumination is not perfectly coherent is not taken into account in the current algorithm. The reconstruction process optimizes the optical field at the object plane while maintaining a perfect agreement with measurement at the sensor plane. The reconstruction contains regularization terms based on sparsity and total variation constraints [3]. The illumination wavelength, effective pixel size (physical pixel size of the camera divided by the magnification of the system) and the out-of-focus distance are the input parameters. The out-of-focus distance can be determined from the reconstruction performed at different axial positions. A focus determination algorithm can be employed, however, we often use a manual selection. In the time-laps data the defocus is determined only once and used for the reconstruction of the whole movie. The reconstruction of a single image from our 10 Mpixel camera takes approximately 3 minutes on our standard desktop computer (Processor Intel(R) Xenon(R) CPU E3-1240 v5 @ 3.50GHz with 32GB RAM and NVIDIA Quadro K2200 graphic card).

#### IMAGE ANALYSIS

**Particle image velocimetry (PIV):** The images were divided into windows of size $28\times28$ $\mu$m$^2$ with 14 $\mu$m overlap. For each window, a velocity value was calculated as follows: for a given time shift (e.g. $\tau$=1 frame), the spatial correlation of each window with its corresponding time-shifted one was computed over 4 consecutive frames and averaged to improve the signal-to-noise ratio. The peak of this average correlation gives an estimate of the displacement $\delta\mathbf{r}(\tau)$. The process was repeated for different values of $\tau$ and the final velocity was deduced from a linear regression of $\delta\mathbf{r}(\tau)$. The final time resolution is 20 min.

#### MODEL AND NUMERICAL DETAILS

**Details of the SPV model**: Each cell is characterized by its position $\mathbf{r}_i$ and shape as determined by the Voronoi tessellation of all cell centre positions and an energy is

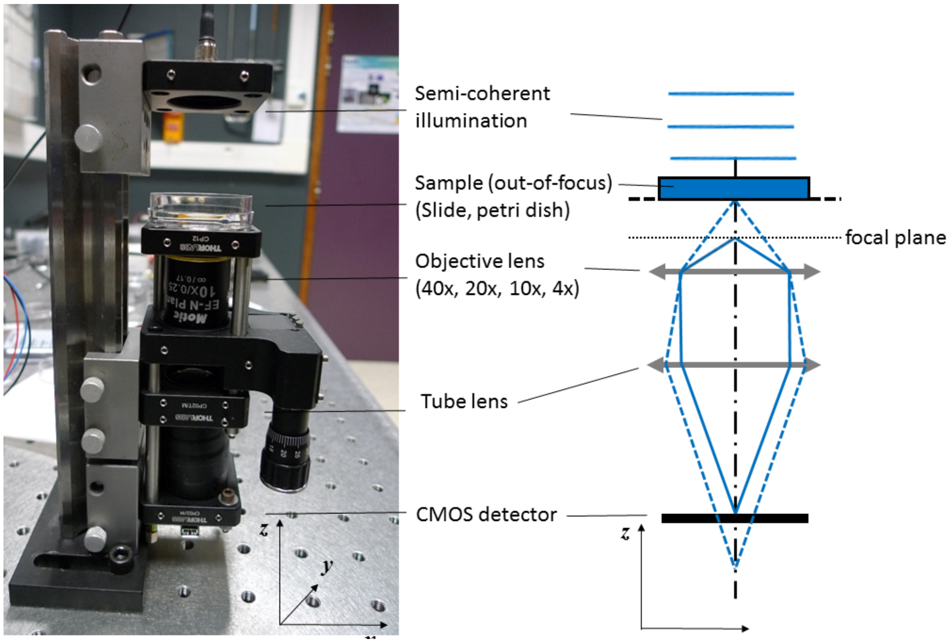


FIG. 1. Setup for defocused imaging is a standard wide-field microscope with semi-coherent illumination.

associated to each configuration of the mesh:

$$E = \sum_{i=1}^{N_{cells}} \frac{K}{2}\left(A_i - A^0\right)^2 + \sum_{i=1}^{N_{cells}} \frac{\Gamma}{2}\left(P_i - P^0\right)^2 \quad (1)$$

$N_{cells}$ is the total number of cells, $A_i$ and $P_i$ are the area and perimeter of the i-th cell, $K$ and $\Gamma$ are the area and perimeter stiffness respectively, identical for all cells. In the overdamped limit, the equation of motion is:

$$\gamma \frac{\partial \mathbf{r}_i}{\partial t} = f_a \mathbf{n}_i + \mathbf{F}_i + \nu_i(t) \quad (2)$$

with $\mathbf{F}_i = -\nabla_{\mathbf{r}_i} E$, the force arising from the tissue energy and from a soft-core repulsion introduced to stabilize the simulations (described as a quadratic potential of stiffness $k_{cc}$ with an interaction radius of half the typical distance between cell centres $a_c$) [4]. $\boldsymbol{\nu}_i(t)$ is an uncorrelated stochastic force and $f_a \mathbf{n}_i$ models the self-propulsive force. The dynamics of the cell polarity $\mathbf{n_i}$, described by an angle $\theta_i$, is described in the main text. Cell division can also be included in this model but does not affect the oscillations, as reported in previous works [5, 6].

**Confinement in SPV:** The SAMoS implementation of the SPV [4, 7] enables for open flexible boundaries which is convenient to model systems with a small number of cells as it is the case in the confined tissues experiments. In SAMoS, the boundaries are imposed through a special type of particles (called "boundary" particles, and denoted $b$ in the parameters list) that form a "boundary line" that delineates between the tissue and its surrounding (described through a boundary line tension $\lambda_b$ and a bending stiffness $\kappa_b$). The confinement is introduced through a rectangular assembly of immobile particles ("wall" of dimensions ($L_X$ , $L_Y$), denoted $w$ in the parameters) that interact through a repulsive potential with the cells, characterized by a stifness $k_{cw}$ and a characteristic length $a_w$. Note that the boundary particles do not represent cells and hence do not interact with the confining layer of immobile particles (interaction potential $k_{bw} = 0$ between the boundary and the wall).

**Simulation methods and parameters:** We integrate the above model using Brownian dynamics [4]. We first prepare the monolayer configurations and oscillations are then studied in steady state with a fixed number of cells. The layer is initialized with a cell number that is fixed by the system size $N_{init} = (L_X - 2)(L_Y - 1)$, and cells are able to divide during a time $T_{growth}$ at a rate $d = d_0(1 - z/\rho_{max})$ (with $z$ the number of neighbours and $\rho_{max}$ a parameter describing the maximum number of neighbours), to achieve a given cell number density. Division is then turned off and the study focuses on times $t > T_{growth}$. The total duration of simulation $T_{run}$ depends on the time period of oscillations $T$ but is usually $T_{run} > 10T$. The parameters of the vertex model ($A^0$, $P^0$, $K$ and $\Gamma$) are chosen such that the monolayer is in a solid-like state ($p^0 = P^0/\sqrt{A^0} = 2.5$) and cells are mostly hexagonal. Note that keeping very low values of $p^0$ is important in order to avoid to have only square cells due to the rectangular confinement constraints, and to prevent shear flow induced by the boundaries, that prevents the

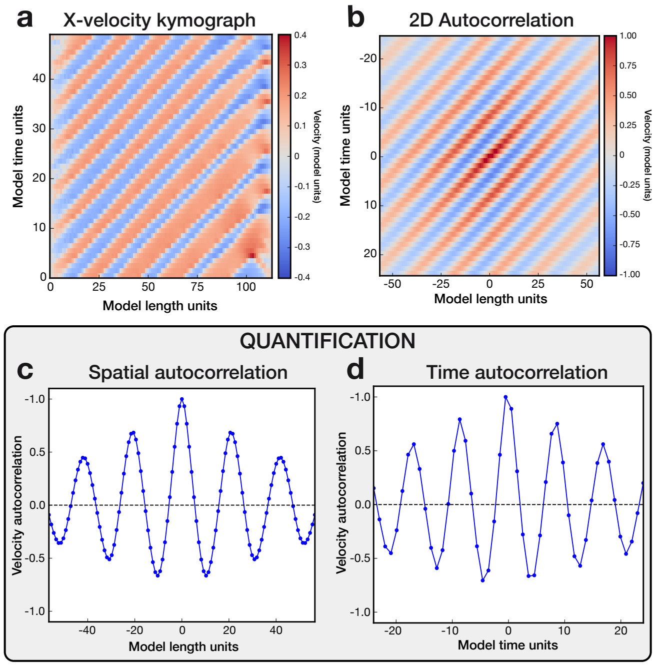


FIG. 2. Illustration of simulation data analysis. a) Kymograph of the x-component of the velocity. b) Autocorrelation of the kymograph. c) Spatial autocorrelation used to extract the wavelength of oscillations. d) Time autocorrelation used to extract the time period of oscillations.

formation of oscillations. In the simulations, the average value of the shape factor $p = \langle P/\sqrt{A} \rangle \simeq 3.85$. The value of self-propulsion velocity is set to $v_0 = 0.2$, but note that changing the value of $v_0$ doesn't affect the features of the oscillations much (it only dictates the amplitude of velocity oscillations). The rotational diffusion coefficient $D_r$ is set to a constant value. The values of all the parameters are listed below.

**Simulation data analysis:** Similarly as for processing experimental data, once in the steady state regime, we average the horizontal component of the centroids velocity along the transverse direction and generate the kymographs of $x$-velocity by displaying this average value as a function of time and space along the horizontal direction $x$. The period and wavelength of oscillations are then extracted from the autocorrelation of the kymograph of x-component of velocity, as shown on Fig. 2.

*
†

[1] https://www.raspberrypi.org/.
[2] J. W. Goodman, *Introduction to Fourier optics* (Roberts and Company Publishers, 2005).
[3] L. Herve, O. Cioni, P. Blandin, F. Navarro, M. Menneteau, T. Bordy, S. Morales, and C. Allier, Biomed. Opt. Express **9**, 5828 (2018).

TABLE I. SPV model, boundary and potential parameters. Raw simulation parameters

| Parameter | Meaning | Value |
|---|---|---|
| $K$ | Area stiffness | 1.0 |
| $\Gamma$ | Perimeter stiffness | 1.0 |
| $A^0$ | Target area | 1.0 |
| $P^0$ | Target perimeter | 2.5 |
| $k_{cc}$ | Repulsive potential strength | 5.0 |
| $a_{cc}$ | Repulsive potential length | 1.0 |
| $k_{cw}$ | Confinement strength (cells) | 5.0 |
| $k_{bw}$ | Confinement strength (boundary) | 0.0 |
| $a_w$ | Confinement potential length | 1.0 |
| $\kappa$ | Boundary bending | 0.1 |
| $\lambda$ | Line tension | 0.1 |

TABLE II. Dynamics parameters

| Parameter | Meaning | Value |
|---|---|---|
| $v_0$ | Self propulsion velocity | 0.2 |
| $\gamma$ | Friction | 1.0 |
| $\gamma_r$ | Orientational friction | 1.0 |
| $\mu$ | Mobility | 1.0 |
| $\mu_r$ | Rotational mobility | 1.0 |
| $\nu_r$ | Rotational noise (rate) | 0.1 |

[4] D. L. Barton, S. Henkes, C. J. Weijer, and R. Sknepnek, PLoS computational biology **13**, e1005569 (2017).
[5] M. Deforet, V. Hakim, H. Yevick, G. Duclos, and P. Silberzan, Nature communications **5**, 3747 (2014).
[6] S. Tlili, E. Gauquelin, B. Li, O. Cardoso, B. Ladoux, H. Delanoë-Ayari, and F. Graner, Royal Society open science **5**, 172421 (2018).
[7] https://github.com/sknepneklab/SAMoS.

TABLE III. Preparation (tissue growth) parameters

| Parameter | Meaning | Value |
|---|---|---|
| $d_0$ | Division rate | 0.5 |
| $\rho_{max}$ | Maximum number of neighbours | 10.0 |
| $T_{growth}$ | Growth time | 4000 |
